# Non-floral scent sources of orchid bees: observations and significance

**DOI:** 10.1101/2024.07.22.604569

**Authors:** Jonas Henske, Bart P. E. De Dijn, Thomas Eltz

## Abstract

Males of the neotropical orchid bees collect environmental volatiles to concoct complex species-specific perfumes that are later used in sexual communication. While perfumes are typically seen as being derived from floral sources, these bees also collect scents from non-floral resources such as decaying wood or tree wounds, even though reports of these sources remain scarce. Here we report observations of male orchid bees collecting scent at 21 different non-floral sources in Central and South America. Male *Eufriesea corusca* that were marked at one of them, a wounded *Protium ravenii* secreting odoriferous sap/resin, returned repeatedly over periods of up to 19 days. Chemical analyses of hind-leg contents suggest that this single non-floral source accounted for a substantial fraction (>50%) of the species-specific perfume. This and other findings strengthen the view that non-floral scent sources play a central role in orchid bee perfume biology. Moreover, at the same *Protium* we also observed female *Euglossa* spp. harvesting resin for nest construction. The collection of substances by both euglossine male and female bees at the same source strengthens the notion that the evolution of male perfume signaling was promoted by a sensory bias for resinous nest construction materials in females.

## 1. INTRODUCTION

The neotropical orchid bees (Euglossini) are renowned for their pivotal role in pollinating numerous tropical plants, particularly hundreds of species of orchids (Vogel, 1966; Dressler, 1968; Janzen, 1971). This is facilitated by their distinctive behavior, first described by Dodson and Frymire (1961), wherein male bees gather exogenous volatiles (Vogel, 1966) and craft intricate perfume blends (Eltz et al., 1999). For volatile uptake, males apply straight-chain lipids that derive from cephalic labial glands to the odoriferous surface dissolving the lipophilic volatiles; this process is comparable to “enfleurage”, the extraction of floral scents in the traditional perfume industry (Whitten et al., 1989). The bees use modified structures on their front- and mid-tarsi to transfer the volatiles to highly specialized hind-leg pouches (Whitten et al., 1989), where individual and species-specific blends accumulate (Eltz et al., 1999; Eltz et al., 2005a; Zimmermann et al., 2009). These blends consist of diverse terpenoids and aromatics (Williams & Whitten, 1983; Eltz et al., 1999; Zimmermann et al., 2009) mixed with labial-gland lipids, whereby the latter are partially recycled and transferred back to the cephalic glands (Eltz et al., 2007). The perfumes are exposed during a characteristic display behavior in the forest understory (Eltz et al., 2005b; Pokorny et al., 2017) and act as intersexual signals for the attraction of receptive females (Henske et al., 2023), arguably reflecting fitness components such as sensory or cognitive acuteness (Henske & Eltz, 2024).

While euglossophilous orchids highly depend on orchid bees for pollination, studies have shown that this mutualism exhibits a one-sided dependency (Ackerman, 1983; Pemberton & Wheeler, 2006), with euglossine-pollinated orchids contributing only a small fraction of the male bees’ perfumes (Ramírez et al., 2011). Time-calibrated co-phylogenies suggested that the evolution of euglossophilous orchids occurred comparatively late, at a time when scent-collecting orchid bees were already largely diversified (Ramírez et al., 2011). Thus, perfume sources other than orchids must have been used at the time the Euglossini originated (Ramírez et al., 2011).

Extant orchid bee males collect volatiles from flowers of a wide array of flowering plant families including, amongst others, Araceae (Williams & Dressler, 1976; Janzen, 1981; Ackerman, 1983), Solanaceae (Williams, 1982; Sazima et al., 1993), Gesneriaceae (Vogel, 1966), Euphorbiaceae (Armbruster & Webster, 1979) and Plantaginaceae (Cappellari et al., 2009). However, the abundance of floral scent sources is highly variable, and orchids in particular are scarce due to low population densities and short-lived flowers (Ackerman, 1983).

Orchid bees have been observed showing scent collection behavior at non-floral sources like decaying wood, rotten and ripe fruits, leaf litter, exposed tree roots, bark wounds, decomposed plant tissue, feces and even walls sprayed with pesticides (Zucchi et al., 1969; Janzen, 1981; Roberts et al., 1982; Ackerman, 1983; Whitten et al., 1989; Whitten et al., 1993; Cappellari & Harter-Marques, 2010). Whitten et al. (1993) hypothesized that these non-floral, potentially more reliable and more abundant sources play a larger role in the concoction of perfumes than previously thought. However, reports of non-floral sources, in particular with accompanying chemical analyses, are still rare. Furthermore, it is largely unknown to what extent orchid bees visit the same source repeatedly over time, and how important single sources can be for the completion of the species-specific blend. Mark-recapture studies are mostly done with artificial scent baits, which offer highly attractive chemicals leading to repeated visits of male orchid bees (Dodson, 1966; Ackerman & Montalvo, 1985; Eltz et al., 1999; Roubik & Hanson, 2004; Pokorny et al., 2013; Pokorny et al., 2015). With the exception of euglossophilous orchids, the entirety of euglossine scent sources remains essentially understudied.

In this study, we report observations of male bees showing scent collection behavior at various non-floral volatile sources. At one source, we marked the bees and observed them visiting the source on repeated occasions. We provide chemical analyses of this and another source to assess the contribution of non-floral sources to the male perfume. Additionally, we report on females gathering resin at a scent source also used by males, which stimulates discussions on the origin of male scent collection behavior.

## 2. METHODS

### 1. Study sites, source recording, marking and sampling of bees

Non-floral scent and resin sources were recorded at seven different localities. The majority of the sources was recorded during the months of March and April in 2019, 2021 and 2022 at the La Gamba Research Station, located in the Puntarenas region within the Golfo Dulce area of Costa Rica. Located adjacent to the Parque Nacional Piedras Blancas in the southern Pacific region of the country, this station is situated in an area characterized by high annual rainfall (5,900 mm) and consistently warm temperatures, with an average of 28°C (Huber et al., 2008). Access to the nearby forest is possible through a trail network originating from the field station, positioned in a small valley. These trails extend into the adjacent forest and follow various paths along a ridge. We walked the trails regularly but not in a standardized manner in 2019, 2021, and sporadically in 2022. Further observations were made in the Munder area of Paramaribo in Suriname, an urban zone bordering rainforest remnants, and in Cultuurtuin in Paramaribo, an old plantation and experimental garden, in 2015 and from 2019 to 2023. Both study sites are situated on the west bank of the Suriname River with average rainfall of 2,000 - 2,250 mm (Nurmohamed et al., 2018). Additional observations were made in the lowland rainforests of Route de Kaw and Montagne des Singes in French Guiana, and around Tiputini Biodiversity Station and near Yasuní National Park in Amazonian Ecuador.

When encountering an orchid bee showing scent or resin collection behavior we marked the site and took photos or videos of the behavior. Bee and plant species were identified whenever possible based on samples or photographs. The sources were usually observed again on following days. In La Gamba, we found one source of sap/resin (S3) to be especially attractive for male as well as female euglossines. Here, we marked male bees, all belonging to the species *Eufriesea corusca*, with numbered plastic tags (Opalith-tags; Holtermann Imkereibedarf, Brockel, Germany) after catching them with a hand-net. After marking, the bees were released. S3 was observed in total on 30 days between 18 March and 14 May 2019. Observation times ranged from approximately 10 min to an hour per day. On 23 March 2019, a sap/resin extract was obtained from the source by wiping a toothpick over the area that was most attractive to male bees. 10 mm of the sample end of the toothpick were immersed in 500 μL of n-hexane (Sigma-Aldrich, St. Louis, MO, USA), as were 10 mm of unused toothpick that served as a control. We extracted hind-legs (also in 500 μL of n-hexane) of two unmarked male *Ef. corusca* caught at the source on 20 March and 23 March 2019. In 2021, we took additional hind-leg extracts of male *Ef. corusca* collecting at S3 (*N*=11) between 29 March and 19 April. Additionally, individual heads (with labial glands) of male *Ef. corusca* were extracted in the same way to obtain control samples of bees’ endogenous compounds. We took two additional sap/resin extracts of the source on 29 March 2021. In 2022, we observed the source on five days. Furthermore, we took a hexane extract from a different source (bark from the fracture of a large fallen branch, S23) and a hind-leg sample from the corresponding male visitor on 2 December 2022.

### 2. Chemical analysis

A HP 5890 II gas chromatograph coupled to a HP 5972 mass spectrometer (Hewlett-Packard, Palo Alto, CA, USA) was used for the chemical analysis of hexane extracts using splitless injection (1 μL). The system was equipped with a DB-5MS column (30 m, 0.25 μm film thickness, 0.25 mm diameter). The temperature of the GC oven was gradually increased from 60 to 300°C at 10°C/min followed by maintaining a constant temperature of 300°C for 15 min. We used ChemStation (Agilent Technologies, Santa Clara, CA, USA) for peak calling, peak integration and spectral library building. Compound identification was conducted by cross-referencing commercial mass spectral libraries (Adams, 2001; NIST/EPA/NIH mass spectral database 2011) in conjunction with our own user libraries, using both mass spectral and retention information. We excluded from downstream numerical analysis of bee perfume all peaks/compounds endogenous to the bees, i.e. cuticular hydrocarbons and all straight-chain lipids (alkanes, alkenes, alcohols, acetates, diacetates, and wax esters) that were also found in the head-extracts. These compounds are known to originate from bees’ cephalic labial glands (Eltz et al., 2007).

For the in-depth study of S3, we visualized the results of the chemical analysis by calculating and plotting the relative abundances of the ten most abundant compounds we found in the hind-leg samples of *Ef. corusca* (*N*=13) and source (*N*=3) extracts. To further test the correspondence of chemical composition, we regressed the relative proportions (percentage peak area) of compounds present in both types of extracts. For this, we used only the samples taken in 2021.

The figures were plotted using ggplot (R; v. 4.2.1). Additionally, we visualized the resemblance between samples by overlaying the total ion chromatograms of one male *Ef. corusca* hind-leg extract and one source-extract using ChemStation.

Furthermore, we analyzed hind-leg extracts of eight individuals of *Ef. corusca* collected in lowland central Panama in April and May of 2009 and 2010. We compared the relative proportions (percentage peak area) of the most abundant compounds between La Gamba and Panama.

## 3. RESULTS

### 1. Observations

We encountered 28 non-floral scent and sap/resin sources attracting either male or female euglossines, or both (summarized in Table 1).

**Table 1:**
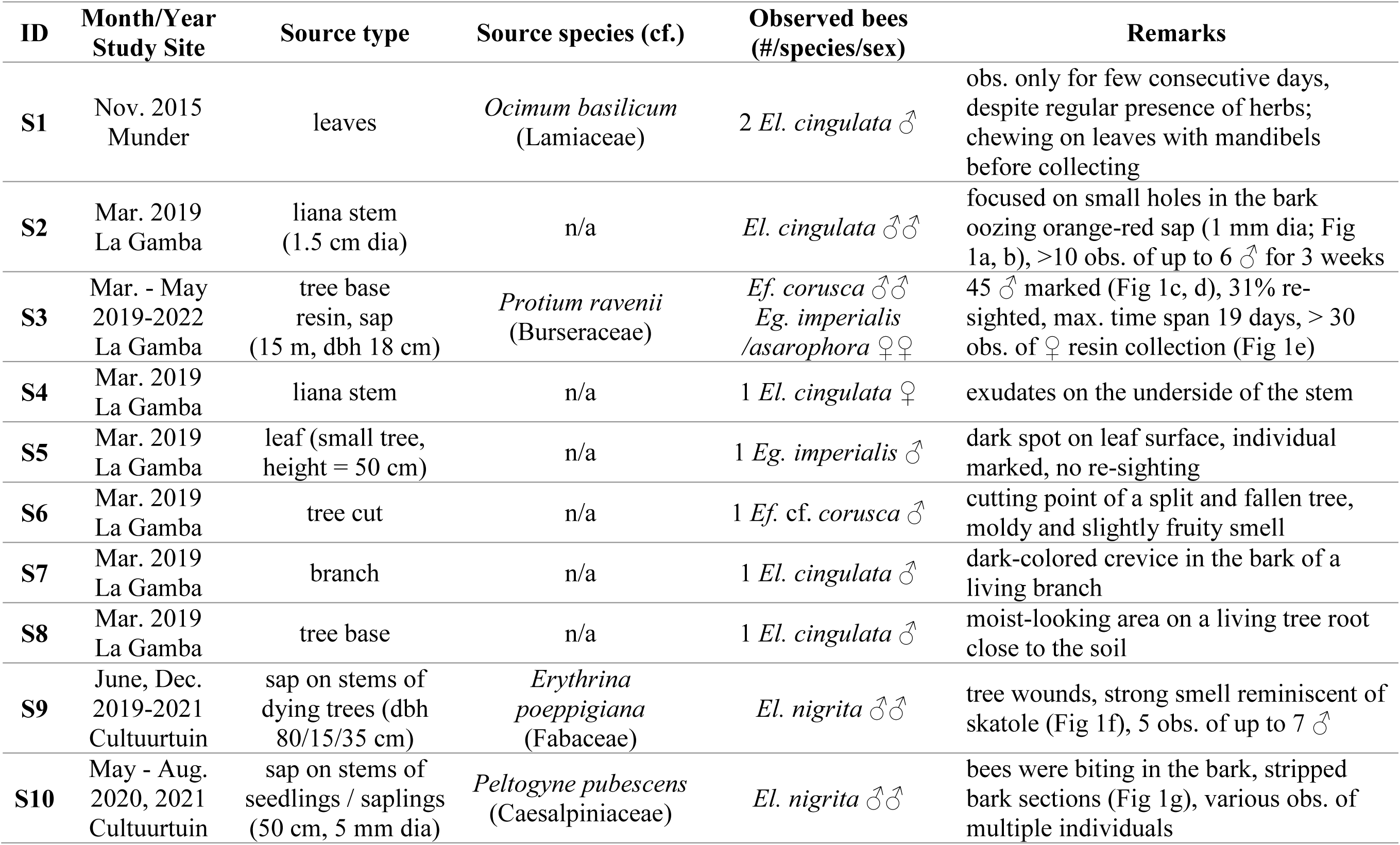
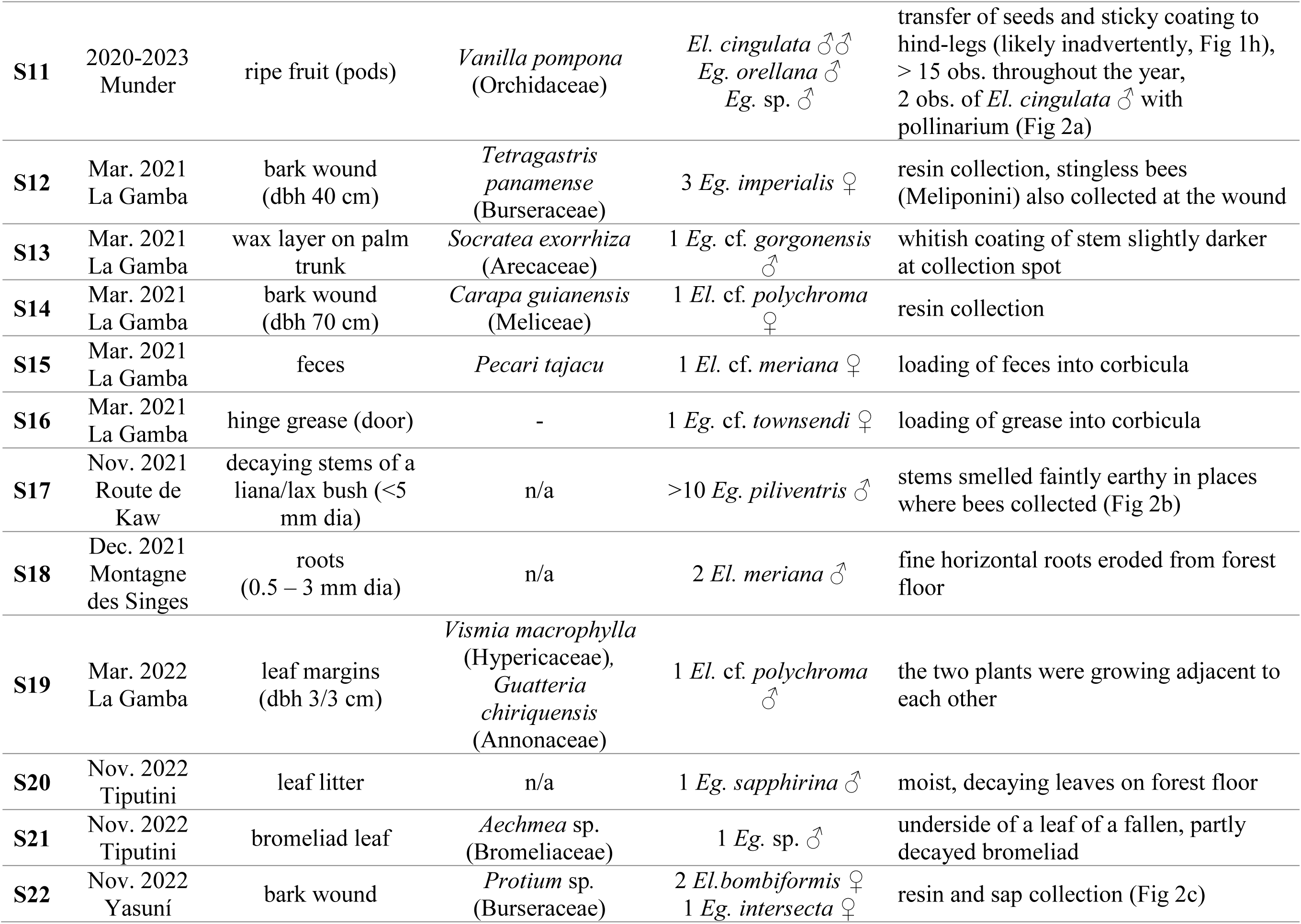
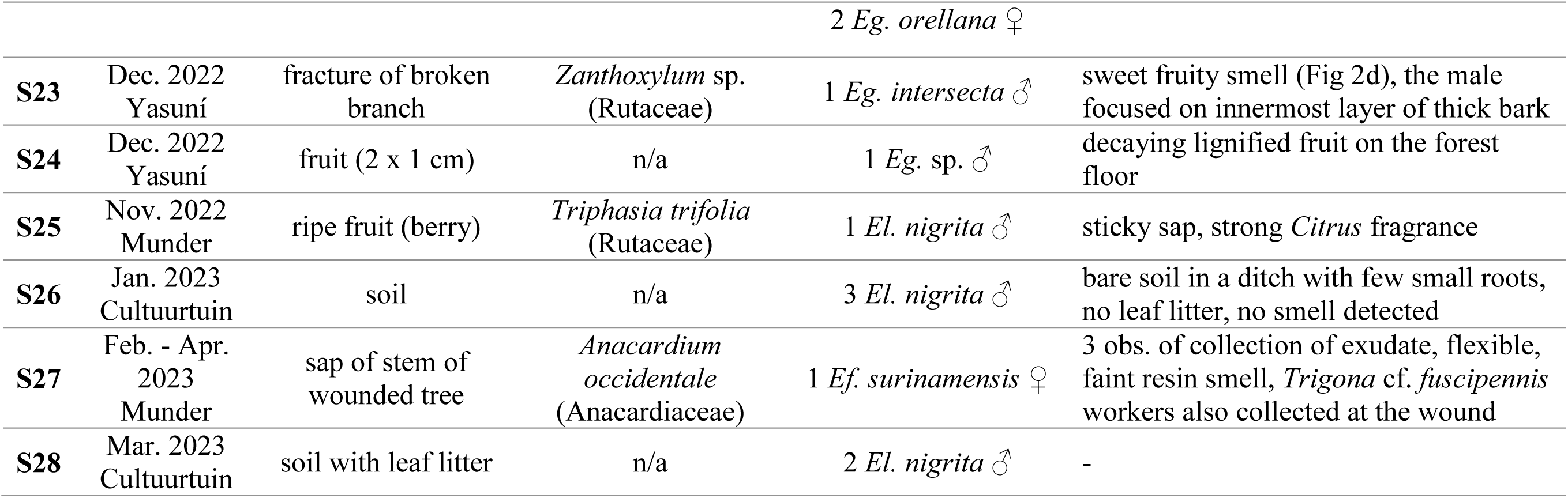
Non-floral scent and resin sources encountered during the course of the study. All observed male bees showed volatile collection behavior as described in Dodson & Frymire, 1961. All observed females collected resin or other specified material into their corbiculae. dbh = diameter at breast height; dia = diameter; obs. = observations; *Eg*. = *Euglossa*; *El*. = *Eulaema*; *Ef*. = *Eufriesea*.

### 2. Chemical analyses

The extracts of S3 contained mostly sesquiterpenes but also some aromatic compounds and other terpenes. The sesquiterpenes germacrene D, δ-cadinene and α-copaene belonged to the five most abundant compounds with an unidentified aromatic compound (molecular weight: 196) and the eudesmane sesquiterpene alcohol junenol being the most abundant compounds across all three samples taken. These five compounds represented on average 69.3% of the total volatile peak area present in the source extracts. The mean total number of compounds found in hind-legs of male *Eufriesea corusca* sampled at S3 was 39 across the two years, ranging from 26 to 48 detected compounds per individual sample. More than half (53.0% +/- 6.9 (m +/- sd)) of the volatiles found in the hind-leg extracts were also present in S3, and those compounds represented 54.2% +/- 8.8 of the total volatile peak area in male *Ef. corusca* hind-legs. Three of the five most abundant compounds found in the bees were also present in the source (junenol: 13.8% peak area, aromatic compound: 10.0%, germacrene D: 7.9%; see Figure 3). Two major compounds were not present in the source (methyl cinnamate: 19.0%, unidentified compound; molecular weight: 235, 10.9%; see Figure 3). The relative proportions of the compounds found in both the source and the perfumes were found to be strongly correlated (see Figure 4). Notably some compounds (e.g. the sesquiterpene β-cubebene; see Figure 3 and Figure S1) were found to be present in the source, but not in the perfumes. These compounds might have become oxidized or have reacted with other compounds present in the hind-legs (Eltz et al., 2019).

To investigate whether *Protium ravenii* was also an important scent source of male *Ef. corusca* in other localities we analyzed hind-leg extracts of eight males sampled in central Panama. These contained a mean total number of 58.3 compounds ranging from 44 to 73 detected compounds. The top five major compounds were the unidentified compound that we already found in the bees sampled at S3 (molecular weight: 235, representing on average 25.7% of the total volatile peak area), bicyclogermacrene (12.7%), atractylone (11.8%), junenol (7.1%) and methyl cinnamate (2.9%). The aromatic compound (molecular weight: 196) we found to be a major compound of S3 was not found in hind-leg extracts, and germacrene D in small amounts (1.6%).

The extract of S23 (bark of *Zanthoxylum* sp.) contained mostly sesquiterpenes and sesquiterpene alcohols. The hind-leg extract of the male *Euglossa intersecta* captured collecting at S23 contained 44 different compounds. Of those, 22 (50%) were also present in S23, representing 60.8% of the total volatile peak area. Of those, (*E*)-caryophyllene was the most abundant compound found in the bee, representing 21.8% of the total volatile peak area in the bee, but only 1.4% in the source extract. Other major sesquiterpenes present in both bee and source were germacrene D (8.7% of peak area), α- and β-selinene (5.1% / 2.9%), α-epi-7-epi-5-eudesmol (3.4%), an unidentified compound (3.0%) and bulnesol (2.0%). The relative proportions of the compounds found in both the source and the perfumes were not correlated (R²=0.013; y=0.21x+2.81).

## 4. DISCUSSION

Our study shows an astonishing diversity and heterogeneity of non-floral scent sources of male orchid bees. Notably, many of the reported scent sources may involve microbial activity. They were either “decaying”, i.e. in a state of microbially assisted decomposition (e.g. S9, S17, S19, S20, S24, S28), or they involved a form of tissue damage that might plausibly result in microbial infection (most other sources). In some cases the original damage was quite obviously inflicted by herbivory, e.g. in S2, a liana where bee activity was restricted to small punctures in the bark. At this point it remains uncertain whether the attractive volatiles were emitted by microbes, fungi or bacteria, or by the plant itself in response to microbial infection (Whitten et al., 1989; Whitten et al., 1993; Schoonenberg et al., 2003; Cappellari & Harter-Marques, 2010). The major attractants of S3, a wounded *Protium ravenii*, were likely plant-derived since *Protium* (Burseraceae) resins, some of them known by the popular name ‘copal’, are known to contain high concentrations of monoterpenes, sesquiterpenes and aromatics (Rüdiger et al., 2007).

Our findings strengthen the view that non-floral scent sources contribute substantially to perfumes of male orchid bees. Previous studies of perfume composition have shown that groups of compounds were highly correlated in relative abundance across individuals, sometimes across species (Eltz et al., 2008; Zimmermann et al., 2009), forming so-called “motifs” within the complex blends. It has been assumed that these motifs are building blocks of perfumes collected together from specific sources, albeit the identity of such sources remained obscure. Most motifs were relatively simple, containing only three or four structurally related compounds (Eltz et al., 2008; Zimmermann et al., 2009), but there is also evidence for larger motifs containing up to eight sesquiterpenes (Darragh et al., 2023). In the present investigation, the relative proportions of compounds in *Protium ravenii* resin (S3) showed a tight correspondence to those of the same compounds in the perfumes of the bees collecting at it, providing the first unambiguous identification of a source motif in euglossine perfumes. One might argue that *Protium ravenii* represents an obligate scent source for male *Eufriesea corusca*. However, conspecific bees from Panama did not share the exact same motif. Whether Panamanian *Ef. corusca* did not collect at *Protium ravenii* or whether the composition of its resin differed from that in Costa Rica remains unknown.

Non-floral scent sources are clearly divers and probably most male orchid bees use non-floral sources at some stage during their volatile foraging. Among the 586 volatile compounds that Ramírez et al. (2011) registered in orchid bee perfumes, 514 had not been reported from scent sources, floral or non-floral. It is likely that many of those compounds originate from non-floral sources. In contrast to flowers, which are often short-lived and restricted to short flowering seasons, non-floral sources may be more persistent in time and thus more available for orchid bees. Especially plant wounds were observed to attract male bees in the present study (S2, S3, S5, S6, S9, S10, S17, S21, S23). In particular, *Protium ravennii* (S3) sap/resin remained attractive for male *Ef. corusca* over two years, eliciting repeated visits by >30 % of the marked individuals over up to 19 days. This is a rare observation of re-visitation of natural scent sources by individual male orchid bees. In fact, we know of only one other study where individually marked euglossine males were sighted repeatedly at a natural fragrance source, flowers of a *Catasetum* orchid (Janzen, 1981). Other reports of individual re-visitation were either inferential (Armbruster & Webster, 1979) or based on artificial, highly concentrated scent baits (e.g. Dodson, 1966; Ackerman & Montalvo, 1985; Eltz et al., 1999; Pokorny et al., 2013; Pokorny et al., 2015; McCravy et al., 2017).

It has only recently been established by controlled experiments that the possession of perfume increases male mating success in orchid bees (Henske et al., 2023). However, it remains entirely unclear what aspect of the perfume, or what compound/motif within the complex blend, is behaviorally relevant. In addition, we do not know the chemical composition of most of the non-floral scents produced by the reported sources. However, at least in some cases our organoleptic observations allow educated guesses at what the bees were seeking: the wounds of *Erythrina poeppigiana* (S9) smelled of skatole, a pungent nitrogen-containing compound, and attracted male *Eulaema nigrita,* which are strongly attracted to baits of synthetic skatole in Suriname and elsewhere (B. De Dijn, pers. obs.). Ripe vanilla pods (S11) smelled of vanillin and attracted *El. cingulata* and *Euglossa orellana*, which are attracted to baits of synthetic vanillin in Suriname (B. De Dijn, pers. obs.). Our observations of orchid bee males removing seeds during volatile collection at pods of *Vanilla pompona* align with previous observations on *V. grandiflora* (Lubinsky et al., 2006), and on *V. planifolia* and *V. odorata* (Karremans et al., 2022). Likely, orchid bee males are at the same time (obligate) pollinators and seed-dispersers of various *Vanilla* species.

Our observations indicate that *Protium ravenii* also represents a long-lasting resin source for female *Euglossa imperialis* and *Eg. asarophora* (Figure 1d). Additionally, we observed females from *Eg. intersecta* and *Eg. orellana* collecting resin from *Protium* sp. in Ecuador (Figure 2c, S22). The utilization of *Protium* (former referred to as *Proteum*) as a resin source for female euglossines is sporadically mentioned in literature (Dressler, 1982; Armbruster, 1984; Rocha-Filho et al., 2012). Notably, these citations are all based on two rather tentative observations, one by Dodson (1966; “[…] probably collected from *Proteum* spp. […]”) and the other by Zucchi et al. (1969; “[…] probably of resin taken from *Proteum* (Burseraceae)”). However, *Protium* is recognized as a resin source for stingless bees (Barth & Da Luz, 2009; Vit et al., in press).

**Figure 1:**
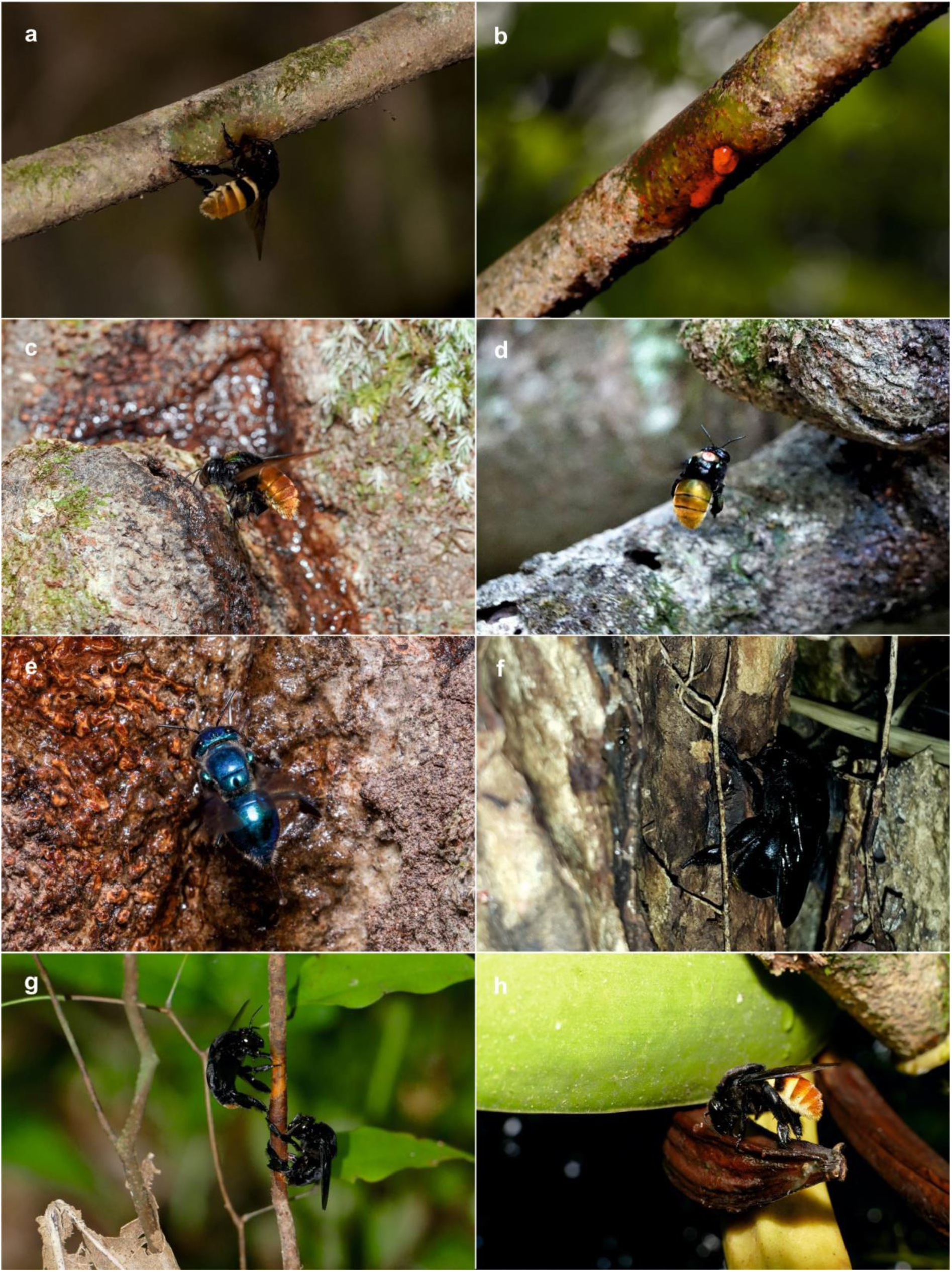
**(a)** *Eulaema cingulata* male showing scent collection behavior at liana (S2). **(b)** Leaking sap at small insect hole (S2). **(c)** *Eufriesea corusca* male showing scent collection behavior at *Protium ravenii* (S3). **(d)** Marked *Eufriesea corusca* male approaching *Protium ravenii* (S3). **(e)** *Euglossa asarophora* female collecting resin at *Protium ravenii* (S3). **(f)** Male *Eulaema nigrita* collecting volatiles on the roots of *Erythrina poeppigiana* (S9). **(g)** *Eulaema nigrita* males biting in the bark of *Peltogyne pubescens* for subsequent volatile collection (S10). **(h)** Male *Eulaema cingulata* collecting volatiles and seeds from ripe pod of *Vanilla pompona* (S11).

**Figure 2:**
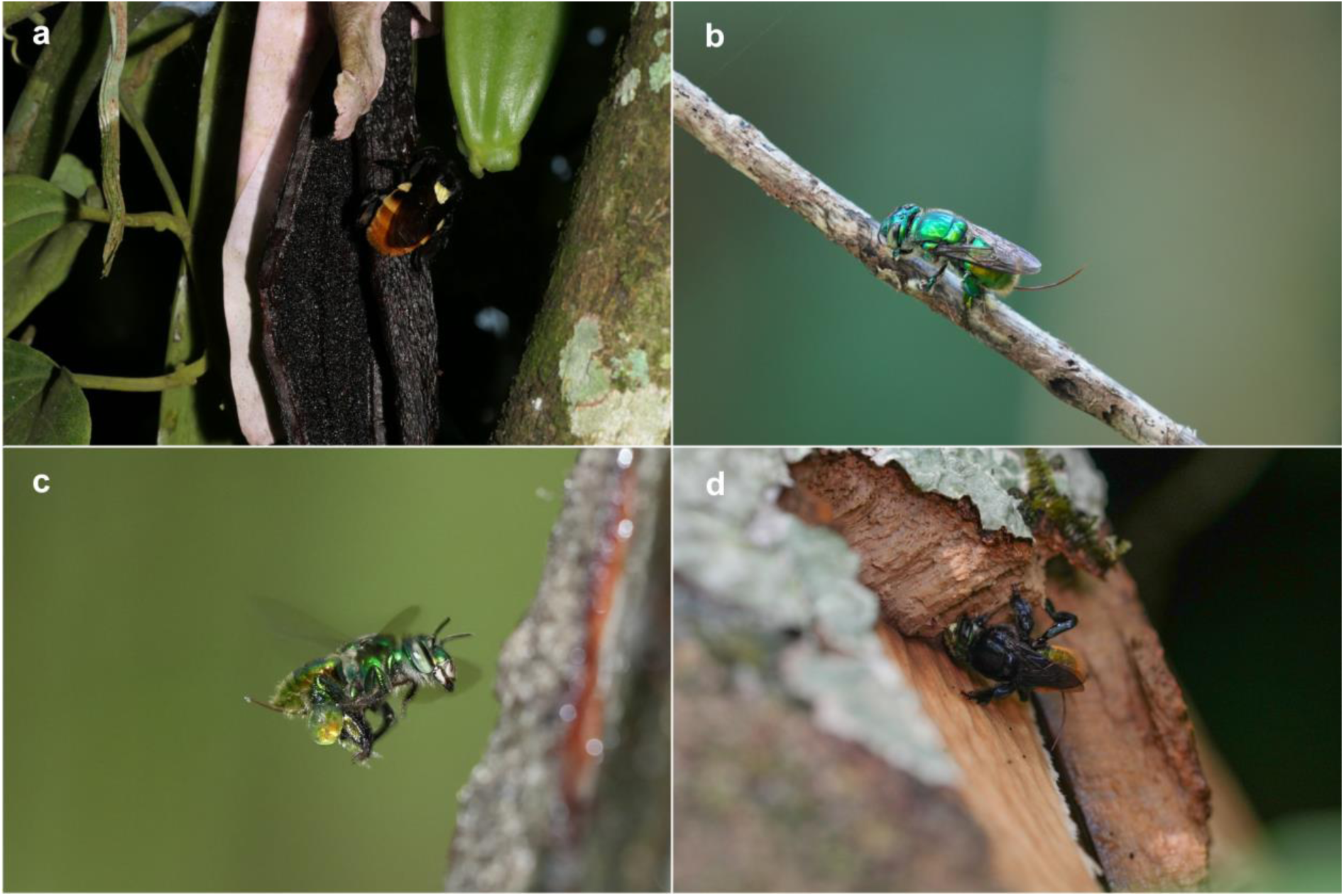
**(a)** Male *Eulaema nigrita* with *Vanilla pompona* pollinarium collecting volatiles from *V. pompona* pods (S11). **(b)** *Euglossa piliventris* male showing scent collection behavior at liana (S17). **(c)** *Euglossa imperialis* female collecting resin on *Protium* sp. (S22). **(d)** *Euglossa intersecta* male showing scent collection behavior on broken branch of *Zanthoxylum* sp. (S23).

**Figure 3:**
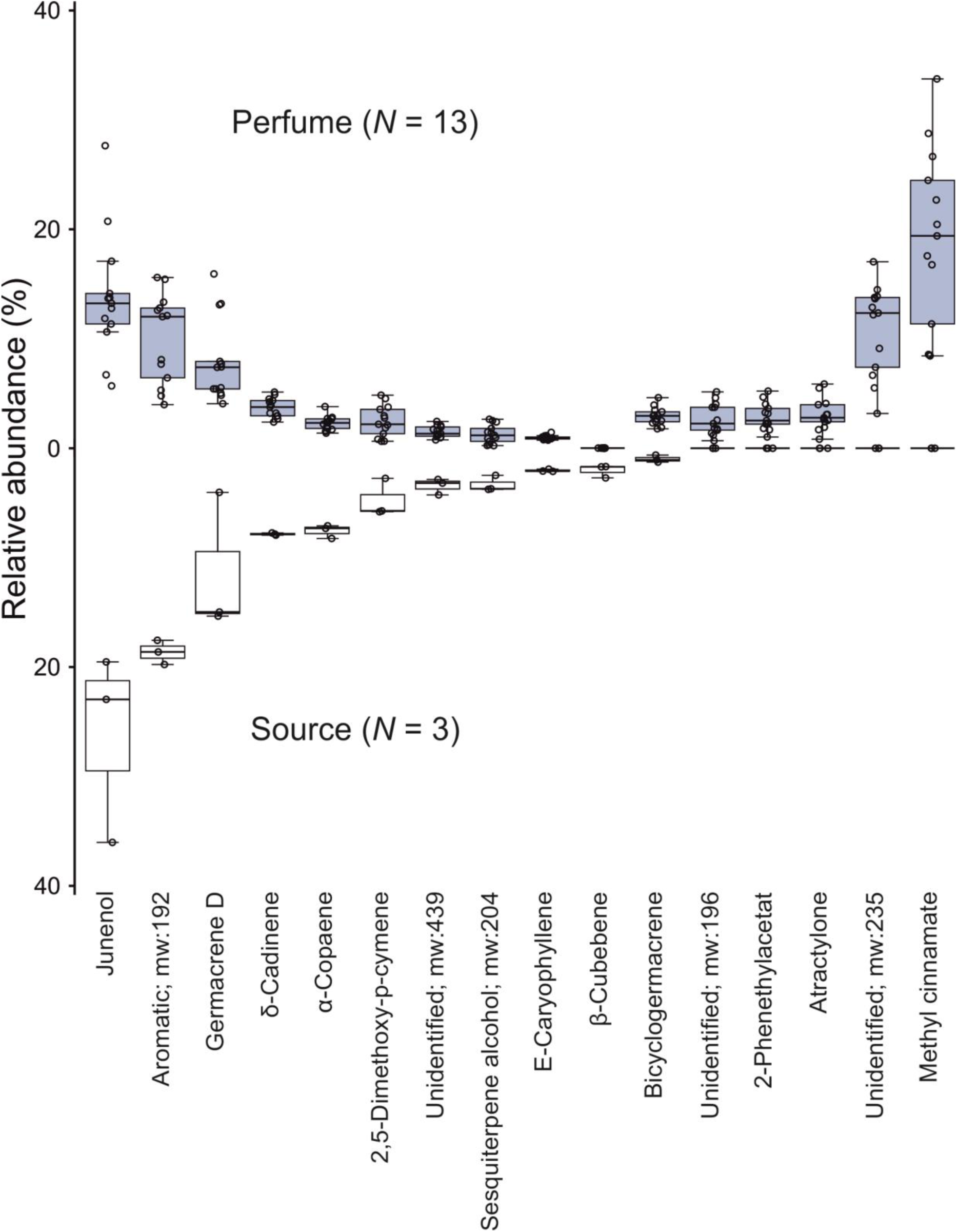
Composition of extracts of resin of *Protium ravenii* (source S3) and hind-legs of the attracted male *Eufriesea corusca.* The relative abundance of the ten most abundant compounds in the perfumes (above) and the source (below) for all samples taken in 2019 and 2021 are shown. Boxplots represent relative abundances (average percentage contribution of total volatile peak area) and show median (center line), upper and lower quartile (box limits), 1.5x interquartile range (whiskers) and individual data points (dots). Compound assignment was based on mass spectra and retention time, mw=molecular weight. See also Figure S1.

**Figure 4:**
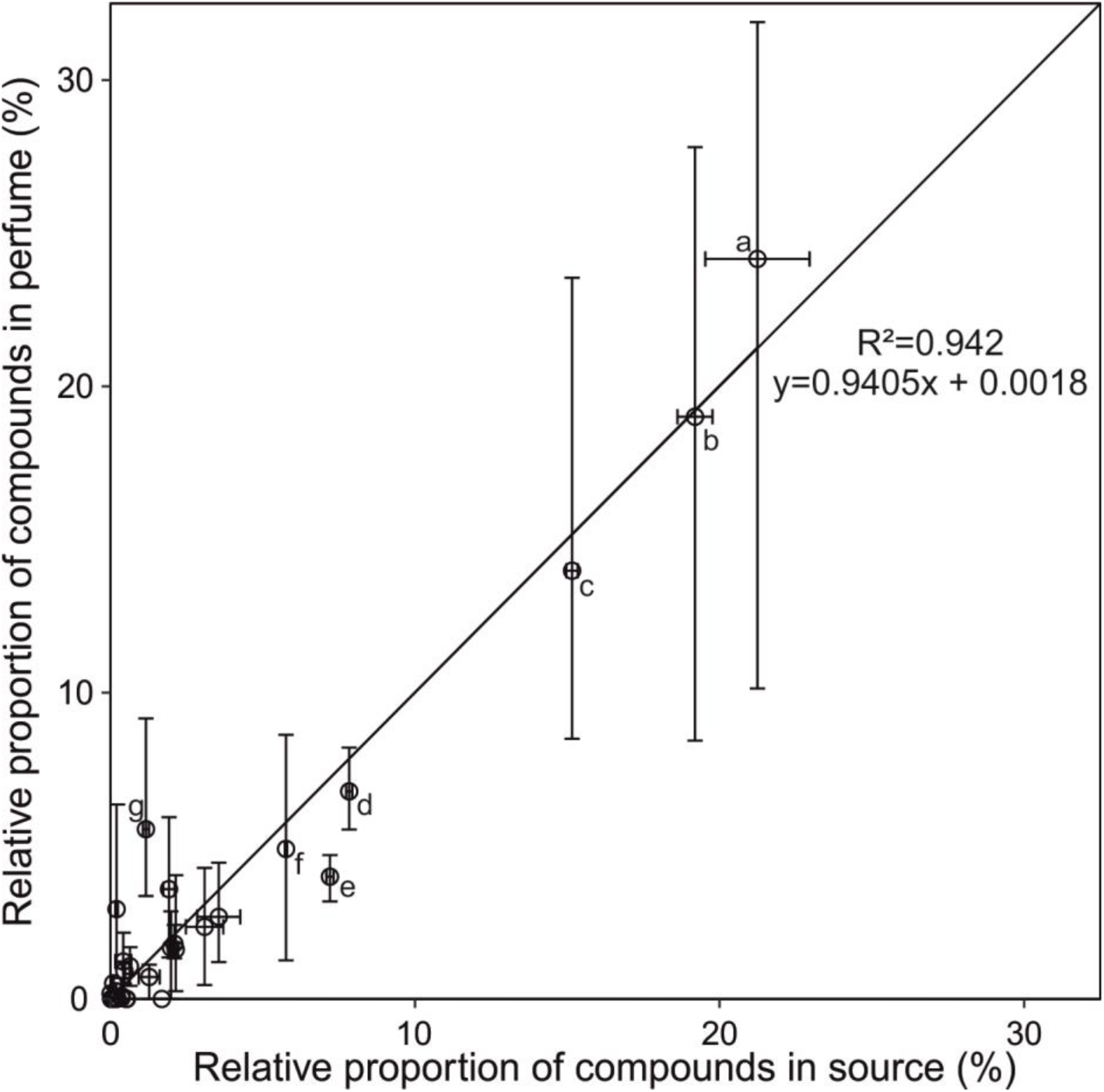
Relative proportions of compounds that overlapped in the source (S3, x-axis) and the perfumes (y-axis) in 2021. Dots indicate mean values of perfume samples (*N*=11) and source samples (*N*=2). Error bars indicate minimum and maximum proportions. Letters represent chemical compounds (assignment based on mass spectra and retention time) and are given when proportion is >5% in source or perfume. **(a)** junenol, **(b)** aromatic; molecular weight: 192, **(c)** germacrene D, **(d)** δ-cadinene, **(e)** α-copaene, **(f)** 2,5-dimethoxy-p-cymene, **(g)** bicyclogermacrene.

Although the females and males observed at S3 did not belong to the same species or even genus, the observations are intriguing and may support the hypothesis raised by Lunau (1992) that “at its outset, the collection of volatiles […] was linked to the female collection of nest material.” The resins of *Protium* contain high amounts of terpenoids, some of which are known to occur in the perfumes of various euglossine species (Rüdiger et al., 2007; Darragh et al., 2023) and also some that are highly attractive when offered in synthetic form (Dodson et al., 1969; Roubik & Hanson, 2004). Thus, our observations support the above-mentioned hypothesis by adding the first observation of parallel scent and resin collection in males and females. At the time of the evolutionary origin of male scent collection behavior, resin already played a major role for nest-building female orchid bees (Cameron, 2004). Accidental smell of resin could have conveyed a mating advantage to males due to a pre-existing sensory bias in females, starting the subsequent co-evolution of male active scent collection and female scent-based mating preferences (see Arnqvist, 2006).

## ACKNOWLEDGMENTS

We thank Werner Huber for the identification of plants. Further, we thank Werner Huber and the entire staff of Tropical Station La Gamba for their constant support. TE wishes to thank Tomi Sugahara, David Romo and Catalina Ulloa from Tiputini Biodiversity Station for making the visits possible. This work was supported by the Studienstiftung des deutschen Volkes (JH), the Wilhelm und Günter Esser Stiftung (JH) and the Deutsche Forschungsgemeinschaft (El 249/11, EL249/13; TE). We thank the Ministerio de Ambiente y Energía (MINAE) and the Sistema Nacional de Áreas de Conservación (SINAC) of Costa Rica (N°SINAC-ACOSA-DT-PI-R-003-2023), as well as the Ministerio del Ambiente, Agua y Transicíon Ecológica of Ecuador (MAE-DNB-CM-2019-0122) for granting permission to conduct this research.

## AUTHORS CONTRIBUTIONS

Conceptualization: TE, JH; Methodology: JH, TE; Investigation: JH, TE, BDD; Formal analysis: JH; Visualization: JH, TE, BDD; Resources: TE; Funding acquisition: JH, TE; Supervision: TE; Writing – original draft: JH; Writing – review & editing: JH, TE, BDD

## DATA AVAILABILITY

The data that support the findings of this study are openly available in figshare and will be publicly available as of the date of publication.

