## Supplemental Figure S1 for "Non-floral scent sources of orchid bees: observations and significance"

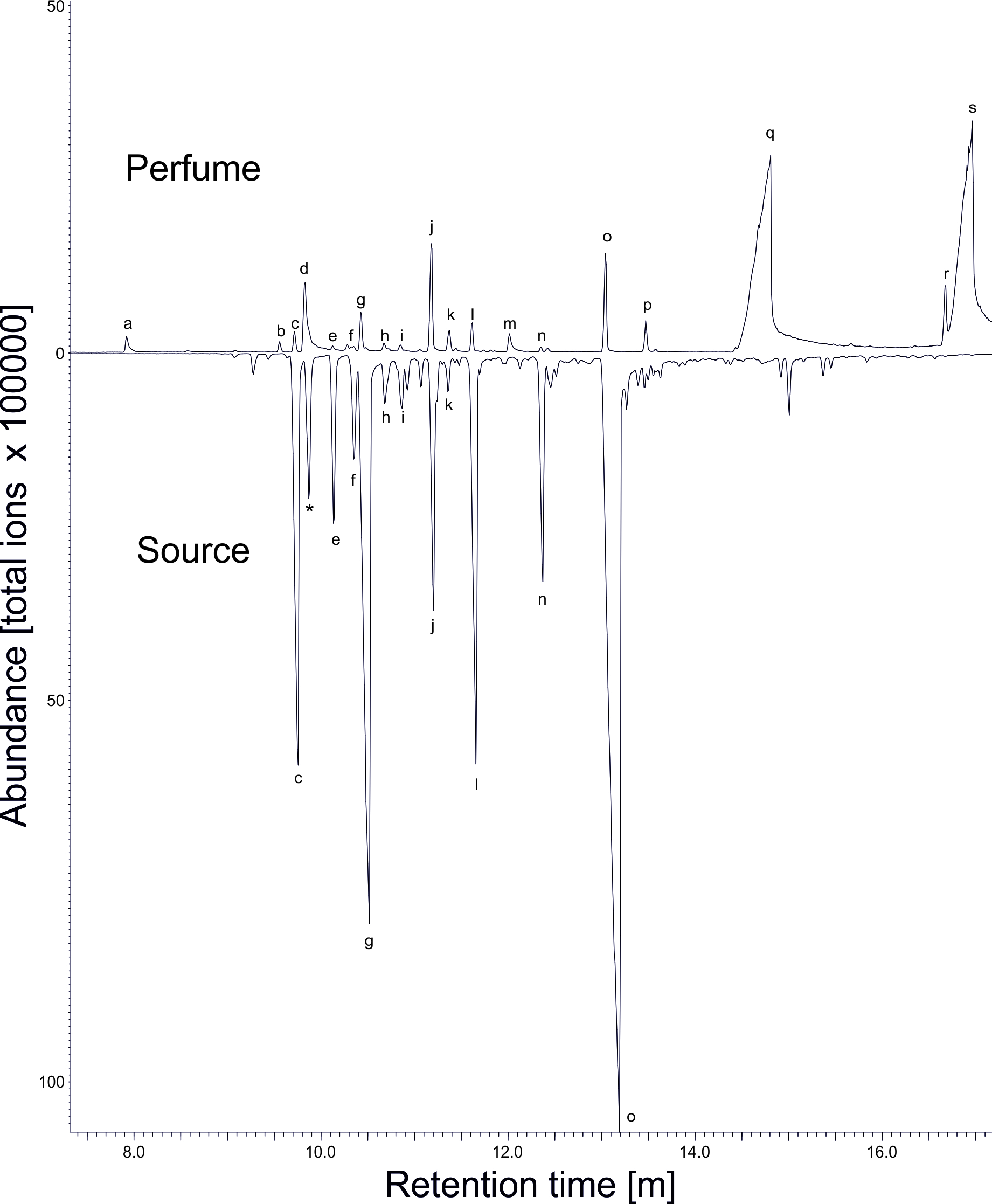

Figure S1: Overlay of total ion chromatograms of extracts of resin of Protium ravenii. (source S3, below) and hind-legs of an attracted male Eufriesea corusca (perfume, above). Letters represent chemical compounds (assignment based on mass spectra and retention time). Compound names are given for perfume compounds but not for compounds that occurred only in the source. **(a)** 2-phenethylacetat, **(b)** unidentified, molecular weight: 202, mass spectra: 159, 131, 105, 145, 91, 133, 117, 202, 77, 119 **(c)** α-copaene, **(d)** methyl cinnamate, **(e)** 2,5-dimethoxy-p-cymene, **(f)** (E)-caryophyllene, **(g)** aromatic, molecular weight: 196, mass spectra: 192, 177, 91, 149, 77, 162, 119, 147, 115, 117 (**h)** sesquiterpene, **(i)** sesquiterpene, **(j)** germacrene D, **(k)** bicyclogermacrene, **(l)** δ-cadinene, **(m)** 1,2-dimethoxy-4-(2-methoxyethenyl)benzene, **(n)** 10-epi-cubebol, **(o)** junenol, **(p)** atractylone, **(q,s)** labial gland lipids, **(r)** unidentified, molecular weight: 235, mass spectra: 145, 219, 105, 91, 55, 79, 107, 93, 77, 234 **(*)** β-cubebene eluted parallel with compound **d** in perfume samples but traces of this sesquiterpene could be found in only one (the shown) hind-leg extract.
